# Communication-Efficient Cluster Scalable Genomics Data Processing Using Apache Arrow Flight

**DOI:** 10.1101/2022.04.01.486780

**Authors:** Tanveer Ahmad, Chengxin Ma, Zaid Al-Ars, H. Peter Hofstee

## Abstract

Current cluster scaled genomics data processing solutions rely on big data frameworks like Apache Spark, Hadoop and HDFS for data scheduling, processing and storage. These frameworks come with additional computation and memory overheads by default. It has been observed that scaling genomics dataset processing beyond 32 nodes is not efficient on such frameworks.

To overcome the inefficiencies of big data frameworks for processing genomics data on clusters, we introduce a low-overhead and highly scalable solution on a SLURM based HPC batch system. This solution uses Apache Arrow as in-memory columnar data format to store genomics data efficiently and Arrow Flight as a network protocol to move and schedule this data across the HPC nodes with low communication overhead.

As a use case, we use NGS short reads DNA sequencing data for pre-processing and variant calling applications. This solution outperforms existing Apache Spark based big data solutions in term of both computation time (2x) and lower communication overhead (more than 20-60% depending on cluster size). Our solution has similar performance to MPI-based HPC solutions, with the added advantage of easy programmability and transparent big data scalability. The whole solution is Python and shell script based, which makes it flexible to update and integrate alternative variant callers. Our solution is publicly available on GitHub at https://github.com/abs-tudelft/time-to-fly-high/genomics.

## 1 INTRODUCTION

Due to massively parallel sequencing methods used in high throughput sequencing (HTS) technologies are making their way from research to the field in a wide range of applications ranging from clinical diagnostics to agriculture research. Next Generation Sequencing (NGS) technologies like Illumina short-reads (a couple of hundred bases), can produce high throughput and higher depth DNA sequence coverage at low cost. Similarly, longer read third generation sequencing technologies are also emerging as a more competitive alternative in terms of cost and throughput with improving accuracy as compared to NGS. They can produce reads of up to hundreds of kilobases (kbps). Depending on the experiment design, the need of sequencing coverage varies [19]. A typical 300x coverage human genome dataset size exceeds 2 TBytes [19]. Processing this amount of data on a single computing machine can take multiple days to complete.

High-throughput sequencing technologies are also enabling cancer diagnoses and treatment beyond histopathology and traditional standard-of-care therapies. Molecular and genomic profiling for patients and tumours at time of diagnosis help in improving diagnostic accuracy, better predict outcome, and personalize therapy [10]. Sequencing coverage influence both the accuracy and sensitivity of such genomics analysis. In pediatric brain-tumor studies [10], more than 200x coverage for the tumor sample, and more than 100x coverage for the normal sample are collected for better focus on the concerned tumors.

In the coming years, as sequencing becomes a normal practice for human health and other types of research, single node compute resources to any organization will not be adequate to fulfill the sequencing requirements. The increased need for data processing will lead to use cluster scaled solutions and outsourcing genomics computations to external private and public cloud services on data centers.

Genonmics data processing pipelines (e.g, short-variants, structural variants and copy-number variants discovery) involve many computational processing steps. Sequence alignment and variant calling are two important steps while intermediate steps like sorting, duplicates removal and base quality score recalibration which use row-based SAM/BAM format to store the outcome of these algorithms on I/Os. Generally genomics data formats (FASTQ/SAM/BAM) permit independent compute and analytic operations on a granular level, i.e., even smaller chunks can be processed without any dependency issues. This eventually helps to run genome analysis algorithms on multiple data chunks in parallel. Halvade [7], which uses the Hadoop MapReduce API, while ADAM [12] and SparkGA2 [13] use the Apache Spark framework and HDFS as a distributed file system are few examples of frameworks which use big data frameworks to scale-up variant calling pipelines on clusters. Because big data scalability requires moving a lot of data between nodes in a big data analytics infrastructure, the current row-based data storage formats and processing row-by-row make these frameworks less efficient for linear scalability and high performance. These solutions use Apache Spark/Hadoop as big data frameworks loosely integrate existing single node pre-processing and variant calling applications. ADAM [12], for example, introduces its own formats, APIs and processing engines. It is built on top of Avro and Parquet for columnar I/O based storage. These solutions come with extra memory overhead and scalability issues.

In order to address these overhead challenges, Apache Arrow in-memory columnar data format in genomics applications has been shown to provide for efficient storage, in-memory analytics and better cache locality exploitation in addition to improved parallel computation [3, 4]. However, limitations in communication overhead remain a challenge, thereby limiting scalability potential of these solutions compared to their custom-made MPI-based HPC alternatives. In this work we establish a case for low-overhead communication using Apache Arrow Flight, enabling efficient scalability of pre-processing and variant calling applications for NGS data on a cluster. This solution leverages the benefits of the Arrow in-memory columnar data format and Arrow Flight wire-speed protocol for shuffling data (between nodes) to sort reads after alignment. The whole workflow is created in Python and Pandas dataframes, which enable computation/analytics like sorting and duplicates removal of NGS data. This solution combines the easy programmability and flexibility of big data pipelines with the high performance and scalability of it HPC-based alternatives. The main contributions of this work are as follows:

- BWA-MEM, a sequence alignment algorithm has been modified to output Arrow in-memory columnar data instead of SAM file. Each BWA-MEM instance on each executor creates 128 Arrow RecordBatches corresponding to chromosomes. This approach stores chromosomes regions level sorted SAM reads.
- Arrow Flight data communication (receiver and transfer protocol) applications have been developed, which communicate with each other to shuffle data through Arrow Flight endpoints.
- On each executor node, Arrow data is converted to Pandas dataframes through PyArrow APIs. A Picard MarkDuplicate compatible algorithm for short-reads duplicates removal is developed in Python.
- The whole variant calling pipeline (alignment, sorting, duplicates removal and variant caller) is managed through SLURM workload manager scripts to use in-memory data for intermediate applications.

In summary, this implementation has following advantages over the existing Apache Spark and MPI based workflows:

- As compared to Apache Spark based frameworks, this approach provides more than 2x speedup, better cluster scalability, less memory footprints, efficient system resource utilization and low communication overhead for data shuffling in intermediate applications.
- When comparing with MPI based solutions, this approach has similar performance for runtime but exhibits better cluster scalability. However, Python ease of programmability and simple Arrow Flight based cluster creation through SLURM or with any other workload manager makes this approach more attractive and suitable for people with little knowledge of HPC systems and performance scalability.

This paper is organized as follows. In Section 2, a brief introduction of the Apache Arrow in-memory data format, the Arrow Flight protocol and the SLURM scheduler is given. Section 3 outlines some big data based pipelines for NGS data processing. Our implementation for both pre-processing and variant calling is described in Section 4, followed by Section 5, where we compare this approach with existing frameworks in both performance and accuracy. Finally we conclude this work in Section 7.

## 2 BACKGROUND

In this section, we introduce genome sequencing technologies, NGS data, pre-processing and variant calling followed by a short discussion on Apache Arrow data format, Arrow Flight communication protocol and SLURM workload manager.

### 2.1 Genome sequencing and NGS data

To analyze an organism DNA for the purpose of understanding and characterizing the unique features it exhibits, the proper order of bases of its DNA should be determined. Different sequencing technologies have been invented for this purpose. Previously widely used Sanger sequencing, next generation sequencing (short reads) from Illumina and the latest third generation sequencing technologies (long reads) from PacBio and Oxford Nanopore are most common technologies. These technologies produce massive amounts of raw genome sequencing data. To understand it and extract useful information about the DNA bases variations in a genome, multiple computational processing steps are necessary to clean and arrange this data for down stream analysis.

### 2.2 Pre-processing and variant calling

In comparative genomics, variant calling analysis reveals deep insights into nucleotide-level organismal differences in some specific traits among populations from an individual genome sequence data. To accomplish this analysis NGS data requires a number of pre-processing steps including sequence alignment, chromosome based coordinate sorting, PolymeraseChain Reaction (PCR) duplicates removal and sometimes base quality score re-calibration. These steps are common in all most every variant calling workflow.

### 2.3 Apache Arrow

Apache Arrow is an in-memory standard columnar data format. Due to the columnar data storage, efficient vectorized data analytics operations and better cache locality can be exploited using this format. Apache Arrow [5] is becoming a standard columnar format for in-memory data analytics. Introduced in 2015, Arrow provides cross-language interoperability and IPC by supporting many languages such as C, C++, C#, Go, Java, JavaScript, MATLAB, Python, R, Ruby, and Rust. Arrow also provides support for heterogeneous platforms in the form of rapids.ai for GP-GPUs and Fletcher for FPGA systems [14]. Apache Arrow is increasingly extending its eco-system by supporting different APIs (e.g., Parquet, Plasma Object Store, Arrow Compute, etc.) and many open-source libraries/tools are integrating Arrow inside them for efficient data manipulation and transfer. For example, TensorFlow has recently introduced the TensorFlow I/O [18] module to support the Arrow data format, the Dremio big data framework is built around the Apache Arrow eco-system, pg2arrow (a utility to query PostgreSQL relational database), turbodbc which supports queries in Arrow format, etc. Arrow stores data in contiguous memory locations to make the most efficient use of CPU cache and vector (SIMD) operations. Moreover, Arrow can efficiently manage big chunks of memory on its own without any interaction with a specific software language run-time, particularly garbage-collected methods. This way, large data sets can be stored outside heaps of virtual machines or interpreters, which are often optimized to work with few short-lived objects, rather than the many large objects used throughout big data processing pipelines. Furthermore, movement or non-functional copies of large data sets across heterogeneous component boundaries are prevented, including changing the form of the data (serialization overhead).

### 2.4 Arrow Flight

Arrow Flight is a submodule in the Apache Arrow project which provides a protocol for transferring bulk Arrow format data across the network. Apache Arrow is also being integrated into Apache Spark for efficient analytics for columnar in-memory data. Arrow Flight [6] provides a high performance, secure, parallel and cross-platform language support (using the Apache Arrow data format) for bulk data transfers particularly for analytics workloads across geographically distributed networks. Using Apache Arrow as standard data format across all languages/frameworks as well as on the wire allows Arrow Flight data (Arrow RecordBatches) to prevent any serialization/deserialization when it crosses process boundaries. As Arrow Flight operates directly on Arrow RecordBatches without accessing data of individual rows as compared to traditional database interfaces, it is able to provide high performance bulk operations. Arrow Flight supports encryption out of the box using gRPC built in TLS/OpenSSL capabilities. A simple user/password authentication scheme is provided out-of-the-box in Arrow Flight and provides extensible authentication handlers for some advanced authentication schemes like Kerberos. A simple Arrow Flight client-server setup in which clients connect and establish connection to a server and preform DoGet() operations is shown in Figure 1. The performance efficiency and throughput of Arrow Flight connection in a remote client-server architecture have been analyzed. The throughput of DoPut() (client send a stream of RecordBatches to the server) and the DoGet() (client receives a stream back from the server) operations is measured and shown in Figure 2. DoPut() throughput increasing from 1.9GB/s to 4.5GB/s while DoGet() achieves 2.5GB/s to 6GB/s throughput with 1 up to 16 Arrow streams in parallel.

**Fig. 1.**
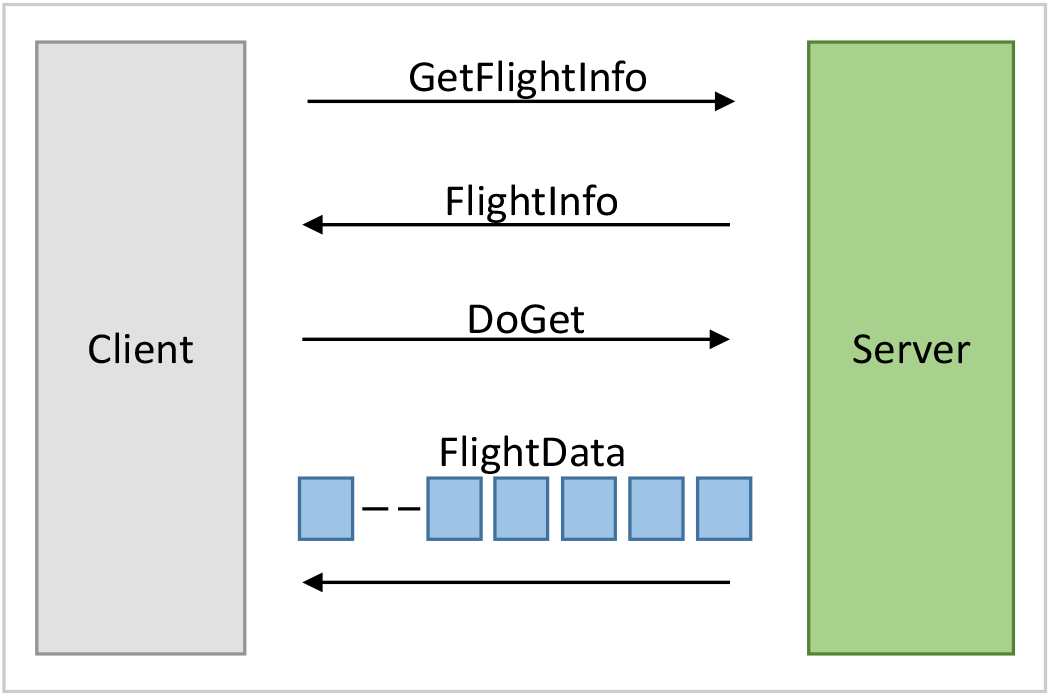
A simple Flight setup might consist of a single server to which clients connect and make DoGet requests.

**Fig. 2.**
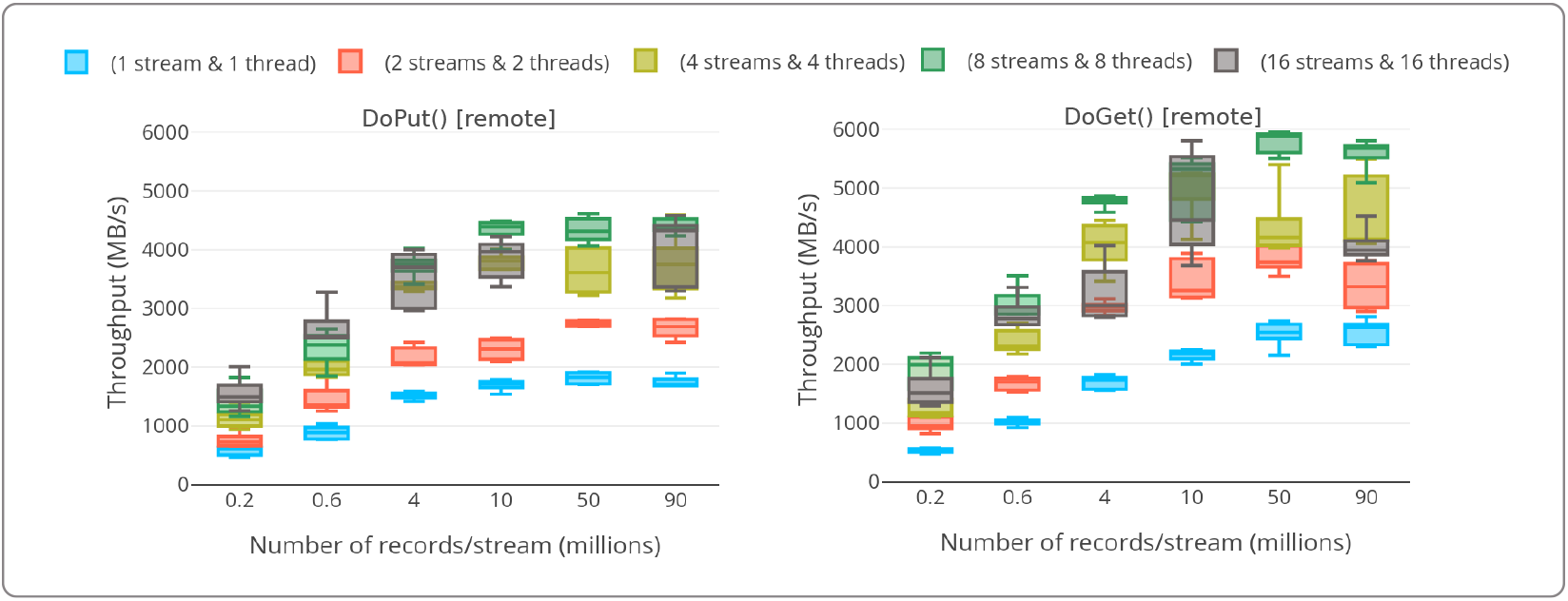
Arrow Flight DoPut() and DoGet() throughput with multiple stream/threads with varying number of records per stream (0.2-90 million) on a remote client-server nodes connected through a Mellanox ConnectX-3 or Connect-IB InfiniBand adapter.

### 2.5 SLURM Scheduler

SLURM is a portable and highly scalable cluster resources management framework. Setting up jobs and resources in SLURM to get bare-metal performance is easy and simple and it also provide both the robustness as well as security needed for HPC applications.

## 3 RELATED WORK

In the past two decades, both high performance computing (HPC) programming models (using MPI) and big data frameworks (like Apache Hadoop and Spark) based solutions have been explored rigorously for genomics applications. Many variant calling workflows and tools have been developed over the past decade, including SparkGA2 [13], ADAM [12], SparkBWA [2], BWASpark [8], etc. Similarly, MPI based parallel versions of the BWA aligner have been developed, such as pBWA [15] as well as QUARTIC, the most recent MPI based BWA (alignment and sorting) [9] algorithm.

## 4 IMPLEMENTATION

While Apache Hadoop and Spark based solutions provide a simple and straightforward method for data parallelization for genomics workflows and particularly somatic/germline variant calling pipelines, still the overheads related to data communication, memory usage and better scalability issues for big clusters remain unsolved for such big data frameworks. We combine the benefits of the Apache Arrow columnar in-memory data format in-conjunction with the high performance wire-speed data transfer protocol, Arrow Flight. SLURM’s managed private cluster is used for distributed and parallel NGS data processing. In the following, the implementation details of pre-processing (alignment, sorting, duplicates removal) applications and variant calling are discussed.

### 4.1 Pre-processing

#### 4.1.1 FASTQ Streaming and BWA-MEM

SeqKit [16] is an efficient multi-threaded command line FASTQ/FASTA data manipulation software. SeqKit runs on a dedicated node and streams out the same number of FASTQ data files as the number of executor nodes available in the cluster in parallel. Each BWA-MEM instance runs on a separate node in a cluster for performance purposes as shown in Figure 3. Each BWA-MEM instance produces in total 128 Arrow RecordBatches in ArrowSAM [4] format. For human genomes, this division into separate chromosome chunks is derived from the ability of load-balanced parallel and independent processing of NGS data on multi-cores and multi-nodes computing systems.

**Fig. 3.**
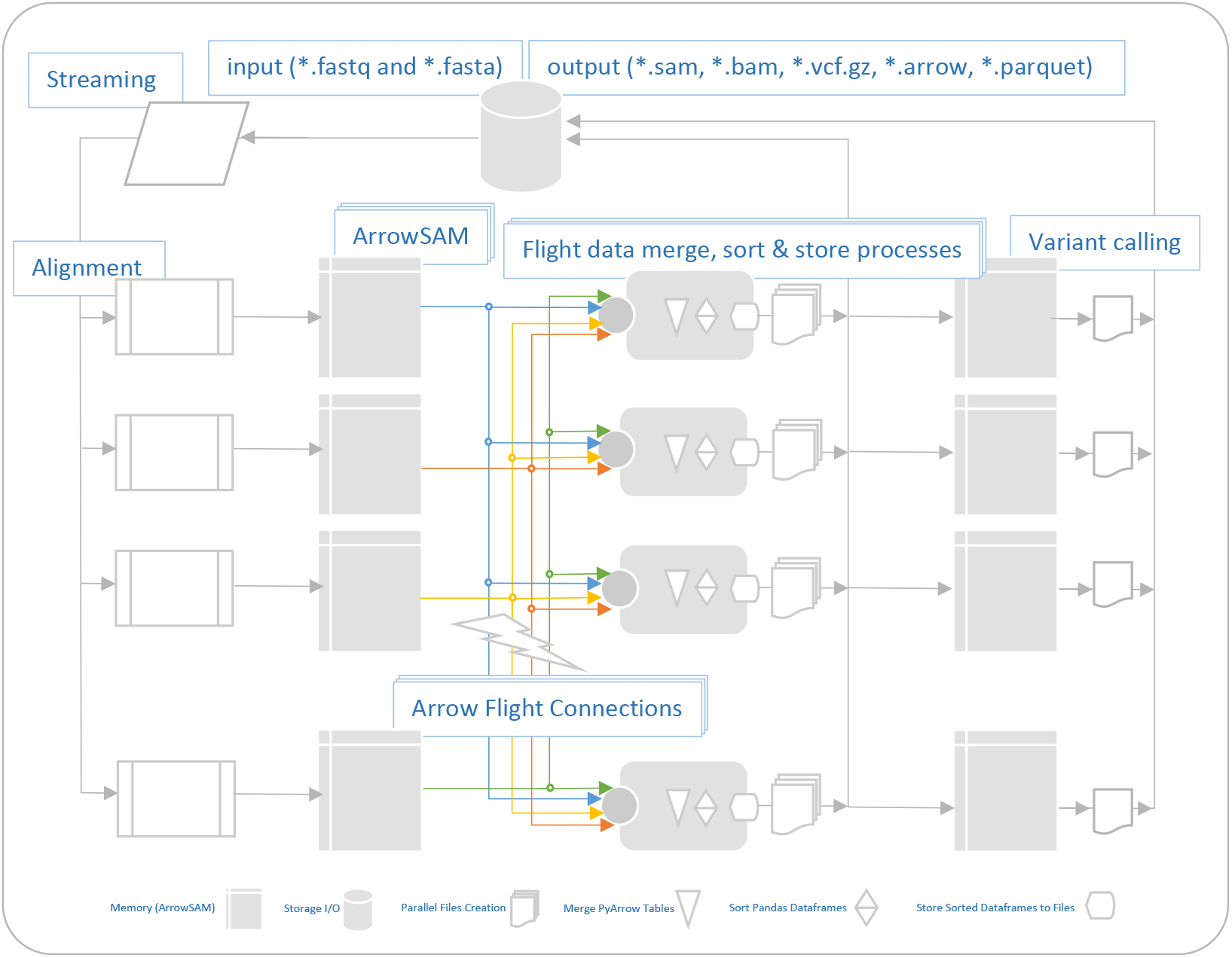
Detailed architectural design of pre-processing and variant calling workflow. Input FASTQ data is being streamed to multiple BWA-MEM instances, which create the ArrowSAM output. Arrow RecordBatches are being transferred/received through Arrow Flight. These Arrow RecordBatches are finally merged, sorted, duplicates removed and resultant output is written on IO, followed by variant calling.

#### 4.1.2 Arrow Flight Data Shuffling

Once the distributed BWA-MEM instances finish the alignment task, Arrow Flight sender and receiver applications on each node start sending/receiving a designated number of Arrow RecordBatches from all the connected nodes through an Arrow Flight connection. Each node sends [128/N] RecordBatches to all available executor nodes, where ‘ N’ is the total number of executor nodes connected through Arrow Flight endpoints.

#### 4.1.3 Arrow Data Merging

Flight receiver application on each node collects total of 128 RecordBatches, coming from its own sender and the sender applications of the rest of the executors. All RecordBatches of specific partitions are then merged through pyarrow APIs and converted to pyarrow tables. The resultant tables are efficiently converted to Pandas dataframes for some further sorting and duplicate removal operations/analytics.

#### 4.1.4 Pandas Dataframe Sorting

All dataframes on each node are sorted by coordinates with the ‘ beginPos’ key in parallel. Sorting on Pandas dataframes with size less than 2GB is efficient.

#### 4.1.5 Pandas Dataframe MarkDuplicate

A Picard “MarkDuplicate” compatible algorithm is developed for duplicate reads removal in pair-end short reads NGS data. The sorted dataframes go through this algorithm by updating the ‘ Flag’ field in case the read in a specific reads bundle is detected as a duplicate.

#### 4.1.6 Intermediate Output

For further downstream variant analysis, any variant caller can be selected in this workflow. All mainstream germline (Strelka2, DeepVariant, Octopus) and somatic (Strelka2, Octopus, NeuSomatic) variant callers use (region-specific) chromosome coordinates like “chr20:10,000,000-10,010,000”. Since this approach outputs data to the I/O with a total of 128 files, this partition is useful when multiple nodes are used for variant calling. The resultant dataframe(s) can be stored on disk in the conventional BAM file format and/or a columnar output file format options like Arrow, Parquet and compressed Parquet for further downstream analysis. The columnar formats are particularly suited for high performance I/O writing/reading.

### 4.2 Variant Calling

Any variant caller which can support region-specific variant calling can be used in this approach. We use DeepVariant a recent and accurate/fast variant caller as a use case to demonstrate the feasibility of using variant caller in this framework.

## 5 EVALUATION

This section evaluates the scalability, throughput and speedup we have achieved for pre-processing of NGS sequencing data in alignment, sorting and marking duplicates stages against the existing frameworks. Here we compare two other existing state-of-the-art cluster scaled pre-processing implementations namely, SparkGA2 [13] and QUARTIC [9].

### 5.1 SparkGA2

SparkGA2 [13] is a Apache Spark based cluster scaled implementation of GATK best practices variant calling pipeline. SparkGA2 starts FASTQ streaming application and initiates multiple BWA instances on Spark executor nodes in parallel. It uses the built-in Scala API in Spark for sorting the aligned reads. As Picard MarkDuplicate algorithm is considered as standard for paired-end reads for duplicates removal, SparkGA2 uses this Picard MarkDuplicate in Spark for distributed processing on cluster.

### 5.2 QUARTIC

QUARTIC (QUick pArallel algoRithms for high-Throughput sequencIng data proCessing) is implemented using MPI. Though this implementation uses I/Os between pre-processing (alignment, sorting and mark duplicate) stages, it still performs better than other Apache Spark based frameworks. These implementations efficiently exploit the multi-cores and multi-nodes parallelization on HPC infrastructure. An MPI wrapper is created for the original BWA-MEM algorithm while using parallel IO and shared memory for alignment. Sorting implements a parallel version of the bitonic sort from scratch in MPI. Their duplicate removal algorithm is based on Picard [1] MarkDuplicate written in MPI.

### 5.3 Experimental Setup

We have performed all the experiments and comparisons on the SurfSara Cartesius [17] HPC cluster (part of the Dutch national supercomputing infrastructure) with each node is a dual socket Intel Xeon server with E5-2680 v4 CPU running at 2.40GHz. Each processor has 14 physical cores with support of 28 hyper-threading jobs. Both processors are connected through Intel QPI (QuickPath Interconnect) and share memory through NUMA (non-uniform memory access) architecture. A total of 192-GBytes of DDR4 DRAM with a maximum of 76.8 GB/s bandwidth is available for whole system. A local storage of 1-TBytes is available on the system. CentOS 7.3 Minimal Server operating system is installed. All nodes are connected through Mellanox ConnectX-3 or Connect-IB InfiniBand adapter.

### 5.4 Data Set

We use Illumina HiSeq generated NA12878 dataset with paired-end reads of WES of human with 30x sequencing coverage. Read length of 100 base-pairs is used for all data. The Human Genome Reference, Build 37 (GRCh37/hg19), is used as a reference genome.

## 6 RESULTS

### 6.1 Performance Evaluation

This section evaluates the performance gains in term of runtime speedups, efficiency and cluster scalability as compared to existing frameworks as well as the Arrow Flight throughput over the network which enables the better overall performance of this approach on clusters.

#### 6.1.1 Runtime Speedup

For pre-processing applications, breakdown of execution time of individual applications is shown in Figure 4. A scalable trend is observed by increasing the number of nodes, also for the communication and data shuffling part in sorting which is explicitly measured here to show a linear decrease in it that is traditionally a bottleneck for scalability on Apache Spark cluster. We also compared the overall runtime of pre-processing applications including BWA, sorting and duplicates removal with the existing state-of-the-art cluster scaled frameworks. Both the Apache Spark based framework (SparkGA2) and MPI implementation (mpiBWA, mpiSORT and mpiMarkDup) have been run on the same cluster with same datasets. Compared to SparkGA2 pre-processing results, more than 2x and 1.5x speedups are achieved, respectively, for all cluster sizes as shown Figure 5. Regarding MPI based comparisons, our approach incurs a marginal overhead for 2 and 4 nodes cluster however increasing the nodes size in cluster to 8 or more nodes, the overall execution time of our approach is decreasing 20%-60%.

**Fig. 4.**
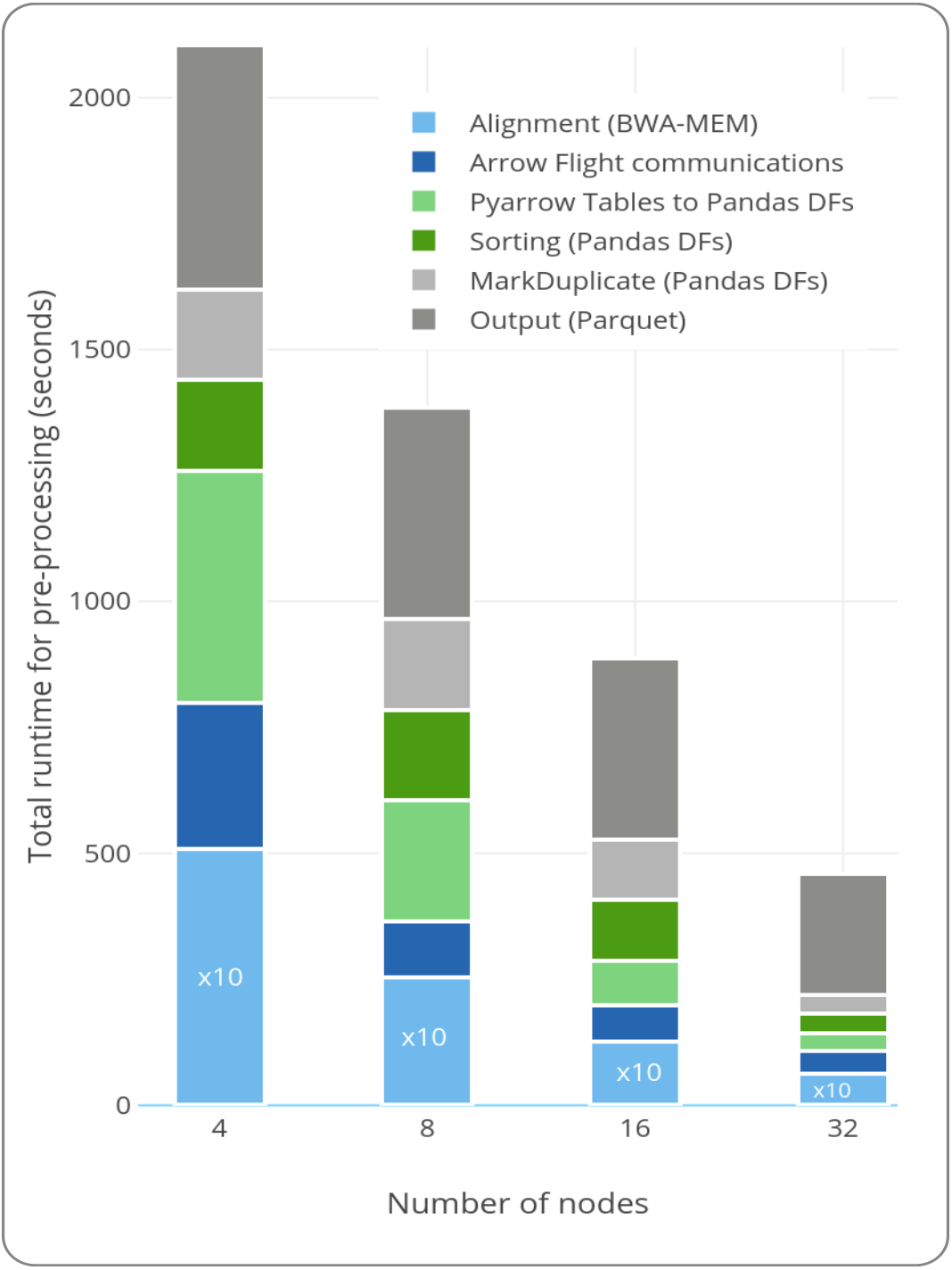
Breakdown of execution time for different pre-processing stages on the HG002 dataset. A scalable trend is observed by increasing the number of nodes, also for the communication part which traditionally is a bottleneck for scalability [13].

**Fig. 5.**
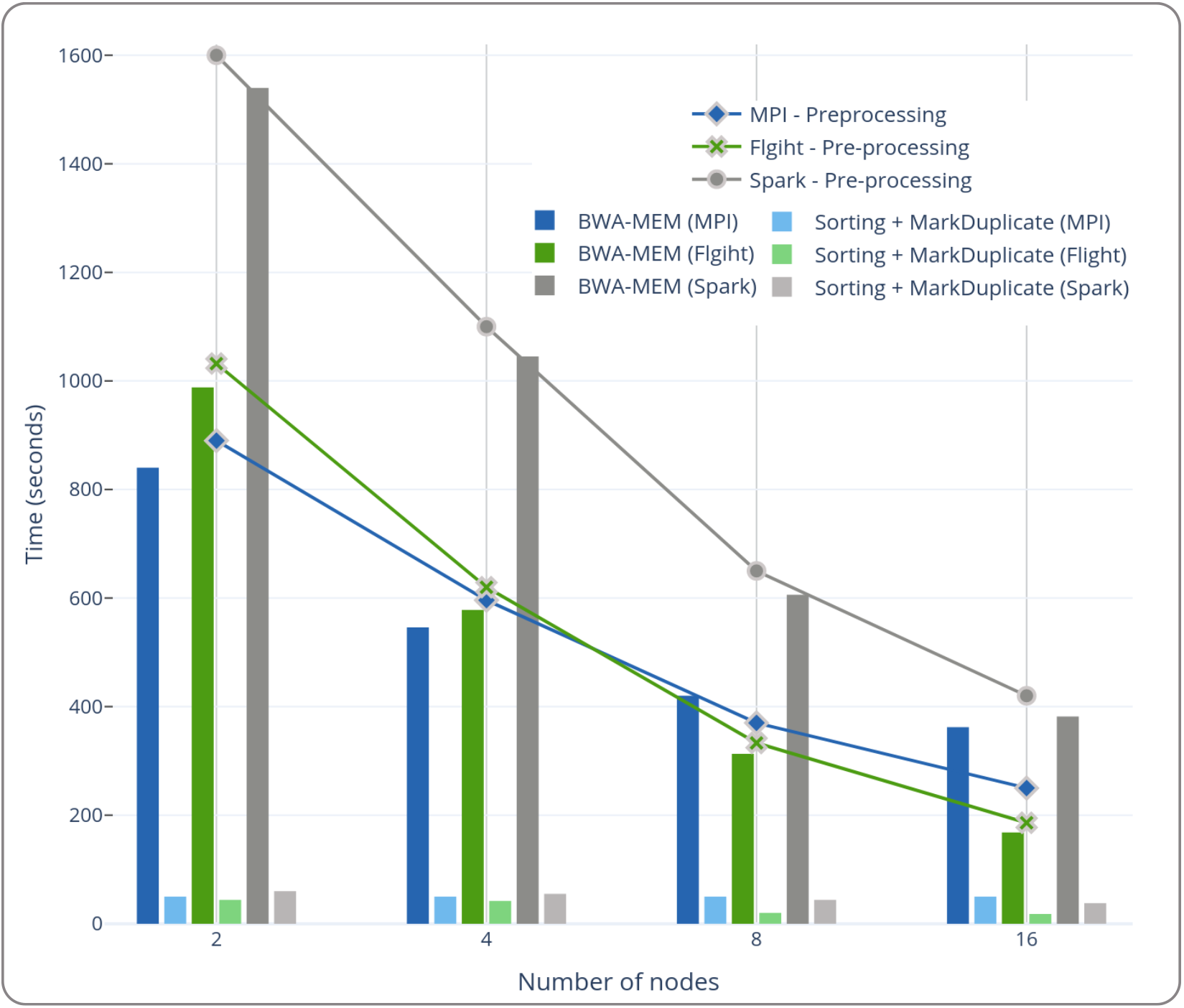
Overall pre-processing runtime performance comparison of different approaches by increasing the number of nodes. Apache Spark (SparkGA2), MPI (QUARTIC) and this approach (Apache Arrow and Arrow Flight) are compared.

#### 6.1.2 Cluster scalability

If the data size remains constant the overall runtime of pre-processing applications can be scaled efficiently by doubling the number of nodes. The total data size also influences the overall performance. As discussed below, Arrow Flight gives better throughput when the Arrow data packet size is big. Normally the Figure 5 does not highlight that linear scalability because the total input data is also being being divided by the factor of nodes being increased.

#### 6.1.3 Memory consumption

With the limited memory available per core on clusters, using the Apache Spark framework incurs additional memory overhead due to its built-in Java and Scala codebase which makes it inefficient to process both computation and memory-bound applications. To prevent this extra memory overhead we replace Apache Spark by SLURM, which is a memory-efficient alternative for cluster environments. As shown in Figure 3 after BWA (alignment) application, data shuffling, merging and sorting and finally duplicates removal is solely being done inside memory. Though this requires almost two folds extra memory of total data size but still this is much less than what is used in Apache Spark based implementations.

#### 6.1.4 Arrow Flight Throughput

It has been observed that a maximum of 4.5GB/s throughput is achievable for DoPut() while DoGet() achieves up to 6GB/s throughput with up to 16 streams in parallel on remote hosts as shown in Figure 2. But increasing the Flight connections in a cluster also effects this throughput. In this implementation, every Arrow Flight connection on each node communicate (sends and receives) with all available Arrow Flight end-points on all the nodes in a cluster. This Arrow Flights connections scenario creates a network congestion but achieves efficient data shuffling. As shown in Figure 6, a maximum 500 MB/s throughput was achievable in a 4 nodes cluster when each node is sending more than total regions files (128) / number of nodes (4) in each iteration to its neighboring nodes at the same time. This also shows that with increasing the Arrow data packet size in each Arrow Flight steams promises much better throughput on even small cluster.

**Fig. 6.**
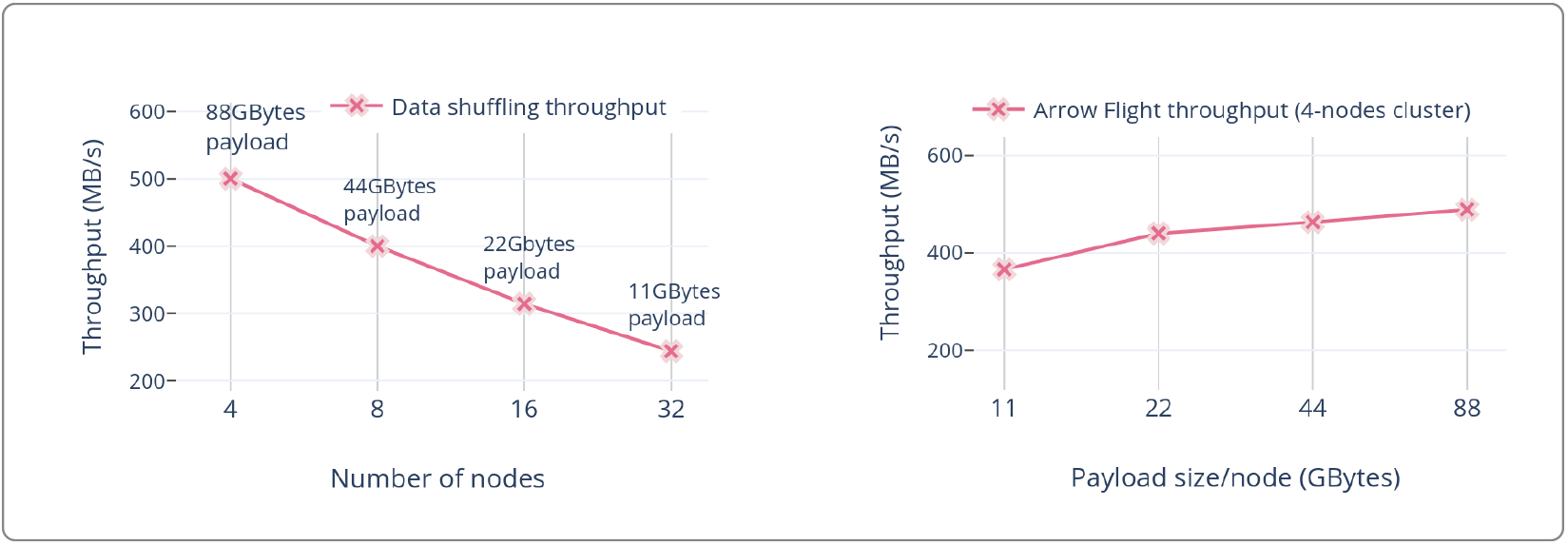
Left figure shows the throughput (on each node) in MB/s with different payload sizes (88-11 GB) with varying number of nodes from 4 to 32, while right figure demonstrates the continued increase in throughput (on each node) when increasing input data size from 11 to 88 GB on a 4 node cluster.

### 6.2 Accuracy

SNP and INDEL variants detection accuracy of DeepVariant variant caller has been compared in a single node and distributed environment. We used HG002 (NA24385 sample with 50x coverage taken from PrecisionFDA challenge V2) dataset to detect SNP and INDEL variants using Deep-Variant (v1.1.0), against GIAB v4.2 benchmark set for HG002 dataset. The GA4GH small variant benchmarking tool hap.py [11] has been used to compare the resulting variants in both methods. Tables 1 and 2 list the accuracy analysis results in terms of recall, precision and F1-score for the single node and distributed approach, respectively.

**Table 1.**
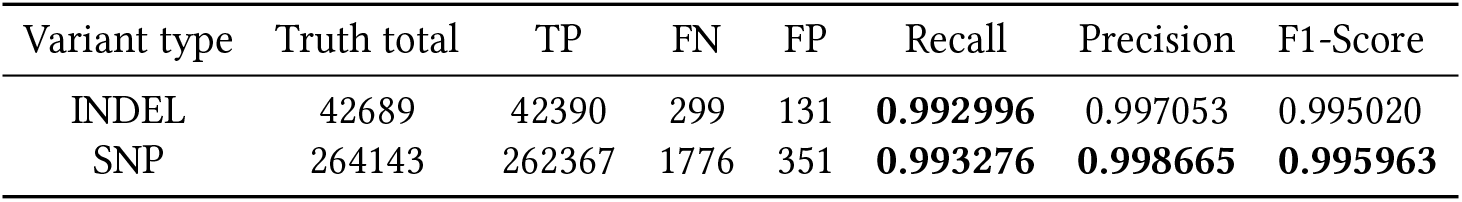
Accuracy evaluation of small variants of HG002 (NA24385 with 50x coverage taken from PrecisionFDA challenge V2 datasets) against GIAB HG002 v4.2 benchmarking set. This table shows the SNP and INDEL results for “Chr1” on a single node (default) run.

| Variant type | Truth total | TP | FN | FP | Recall | Precision | F1-Score |
| --- | --- | --- | --- | --- | --- | --- | --- |
| INDEL | 42689 | 42390 | 299 | 131 | **0.992996** | 0.997053 | 0.995020 |
| SNP | 264143 | 262367 | 1776 | 351 | **0.993276** | **0.998665** | **0.995963** |

**Table 2.** Accuracy evaluation of small variants of HG002 (NA24385 with 50x coverage taken from PrecisionFDA challenge V2 datasets) against GIAB HG002 v4.2 benchmarking set.

| Variant type | Truth total | TP | FN | FP | Recall | Precision | F1-Score |
| --- | --- | --- | --- | --- | --- | --- | --- |
| INDEL | 42689 | 42390 | 299 | 127 | **0.992996** | **0.997142** | **0.995065** |
| SNP | 264143 | 262365 | 1778 | 355 | 0.993269 | 0.998649 | 0.995952 |

## 7 CONCLUSION

This work demonstrates the efficient usage of the Apache Arrow data format and Arrow Flight communication protocol to ensure low-latency communication of genomics data in a cluster environment. Arrow Flight allows for effective scalability of genomics pipelines on large clusters, while eliminating communication time as a scalability bottleneck.

Almost all existing frameworks for processing genomics data are built around big data frameworks like Apache Hadoop and Apache Spark, which does not benefit from columnar in-memory data processing on vector units nor exploit the caches locality efficiently. These frameworks also cost extra memory overheads. Our solution uses the SLURM workload manager as an application handler and data scheduler to replace Apache Spark framework or MPI based implementation of genomics applications. Our approach allows to process more columnar data in-memory without worrying about the extra memory costs. Using SeqKit to create chunks and streaming the resultant FASTQ input to BWA instances eliminates the additional processing time. We have shown that BWA is being scaled almost linearly while initiating only one instance per node. Through this approach we are also able to achieve 1.5x and 2x speedup over existing state of the art frameworks like SparkGA2. The performance comparisons of this approach with MPI based implementation gives similar run-times with better clusters scalability and applications flexibility. Integrating Arrow Flight microservices into existing data transfer and analytics frameworks (Apache Spark, TensorFlow, XGBoost, etc.,) for distributed and scalable processing exhibits both parallel and high throughput data transfer and compute capabilities. Also Arrow Flight based distributed Apache Arrow data scheduling, compute and query services like DataFusion and Ballista present applications for this purpose.

## Notes

https://github.com/abs-tudelft/time-to-fly-high/tree/main/genomics

### Competing Interest Statement

The authors have declared no competing interest.

